# MUTYH cancer-associated variants within the interdomain connector differentially impact glycosylase activity and cellular DNA repair

**DOI:** 10.64898/2026.03.03.709415

**Authors:** Cindy Khuu, Melody Malek, Savannah G. Conlon, Geoffrey P. Wadey, Carlos H. Trasviña-Arenas, Sheila S. David

**Affiliations:** Biochemistry, Molecular, Cellular and Developmental Biology Graduate Group, University of California, Davis, CA 95616; Chemistry and Chemical Biology Graduate Group, University of California, Davis, CA 95616; Department of Chemistry, University of California Davis, Davis CA 95616; Research Center on Aging, Center for Research and Advanced Studies (CINVESTAV), Mexico City, Mexico

**Author notes:** Corresponding Author: Sheila S. David, University of California, Davis, One Shields Avenue, Davis CA 95616.

**Keywords:** Base Excision Repair (BER), MUTYH, DNA glycosylase, 8-oxoguanine, MUTYH-associated polyposis, DNA repair

## Abstract

The base excision repair (BER) glycosylase MUTYH initiates repair of 8-oxo-7,8-dihydroguanine (OG): adenine (A) mispairs to prevent G to T transversion mutations. Inherited biallelic mutations in *MUTYH* are correlated with the cancer pre-disposition syndrome *MUTYH*-associated polyposis (MAP) and contribute to an increased lifetime risk of colorectal cancer. Over 1,000 germline and somatic *MUTYH* variants have been reported that are associated with MAP and other cancers, but for most the functional impact is unknown. Herein, we examined a subset of cancer-associated variants (CAVs) localized in the interdomain connector (IDC), which links the N-terminal adenine excision and C-terminal OG recognition domains via its zinc linchpin motif and serves as a hub for downstream repair interactions. *In vitro* assays measuring glycosylase activity, lesion affinity, and AP endonuclease stimulation revealed no substantial defects relative to wild-type MUTYH. In contrast, a newly optimized mammalian cell assay revealed some IDC variants exhibit reduced repair. These results suggest that some variants disrupt steps downstream of adenine excision, whereas others impair lesion recognition and base excision. This work underscores the value of independent functional assays for accurately assessing variant dysfunction and classification. Analysis of *MUTYH* variants highlights the complexity of the roles of MUTYH in preserving genomic integrity.

## Introduction

MUTYH-associated polyposis (MAP) is a cancer predisposition condition associated with inherited biallelic mutations in *MUTYH*.[1] Patients with MAP are at significant risk for colorectal cancer due to inactivating mutations in the *adenomatous polyposis coli (APC)* gene, a tumor suppressor gene that affects the canonical Wnt signaling pathway to control gene transcription, cell polarity, and motility.[2] The mutation signatures associated with MAP are G:C to T:A transversions that result from compromised ability of the base excision repair (BER) glycosylase MUTYH to remove A from 8-oxoguanine (OG):A mismatches formed during DNA replication (Figure 1).[3,4]

**Figure 1:**
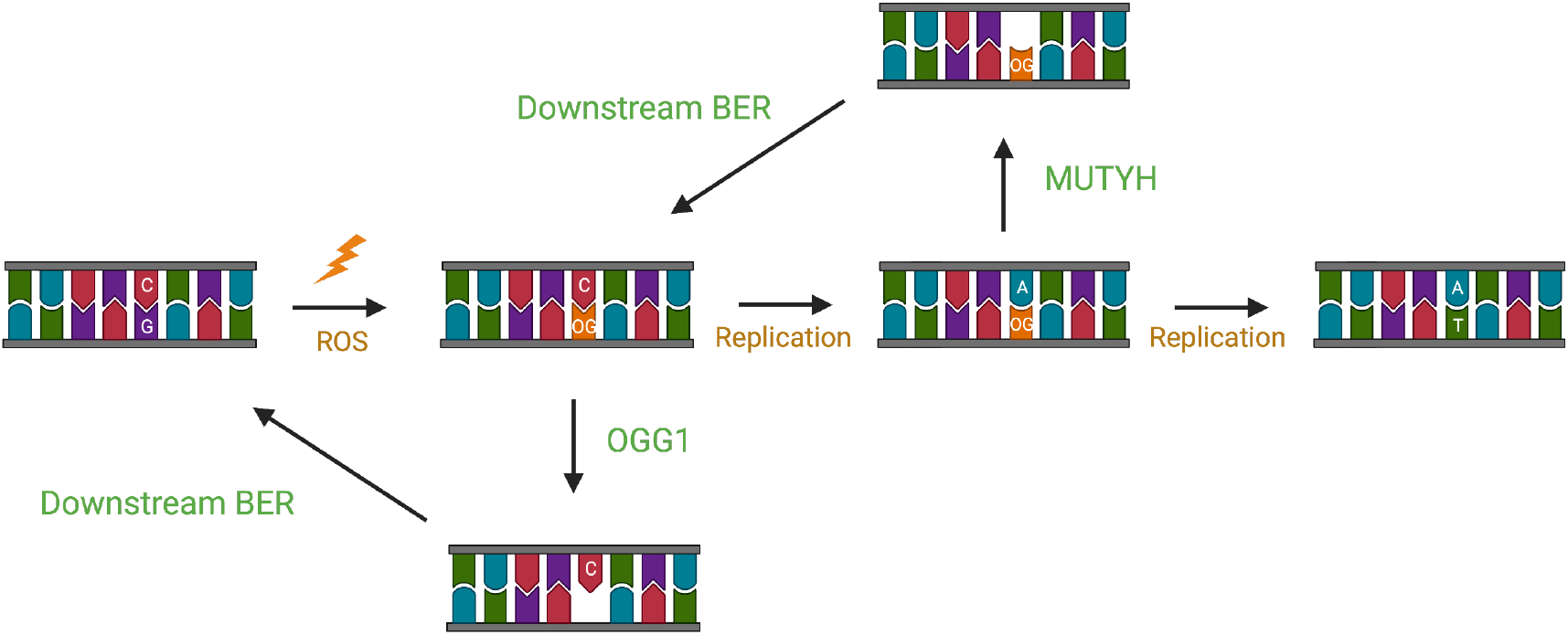
GO repair pathway The base excision repair (BER) glycosylases OGG1 and MUTYH excise aberrant bases in response to guanine oxidation (GO). OGG1 catalyzes the N-glycosidic bond cleavage of OG, creating an abasic site that is further repaired by downstream enzymes. MUTYH catalyzes the N-glycosidic bond cleavage of A incorrectly inserted across OG, creating an abasic site; further processing by downstream BER and OGG1 restores a G:C base pair (bp). Failure to capture OG:A bps results in G:C to T:A transversion mutations.

There are over 1000 single nucleotide variants (SNVs) identified in *MUTYH*, many associated with colorectal and other cancers, including breast, ovarian, skin, and pancreatic.[1–3] Unfortunately, the impact of most of these variants, some that are quite rare, on polyposis and cancer susceptibility are unclear.[5,6] The plethora of mutations detected in *MUTYH* are distributed throughout the protein, and are anticipated to alter different aspects of MUTYH function to varying extents. While the founding MAP variants Y179C and G396D are functionally and structurally well-characterized and have a strong cancer association based on genetics, this is not the case with most variants.[1,5,6] The dysfunction of variants located in functional domains of MUTYH may often be predicted based on structural and biochemical information.[7–10] However, many variants are in regions of the protein that may not alter the base excision chemistry of MUTYH, making functional predictions challenging.

A surprisingly large number of MUTYH variants are localized within the interdomain connector (IDC) that connects the catalytic N-terminal and OG recognition C-terminal domains. In bacterial MutYs, the IDC traverses the DNA major groove to connect the two domains and exhibits minimal secondary or tertiary structure suggesting that it functions primarily as a linker.[11] In contrast, the IDC of human MUTYH is considerably longer (Figure 2).[12,13] Furthermore, we have previously demonstrated that the extended IDC domain in mammalian homologs provides an additional CysX_5_CysX_2_Cys motif that coordinates a Zn ion, which we have referred to as the Zinc Linchpin Motif for its role in coordinating engagement of the functional domains on the OG:A substrate.[14] It has been difficult to gain insight into the IDC using crystallography due to its highly disordered nature. Structures of mammalian MUTYH proteins lack significant electron density for the IDC, and the human protein structure lacked the Zn(II) ion.[12,13] The IDC has been implicated in mediating interactions with several proteins involved in DNA repair and damage signaling, namely AP endonuclease (APE1), HUS1 of the 9-1-1 DNA damage sensing complex, and the stress responsive enzyme SIRT6.[15–18]

**Figure 2:**
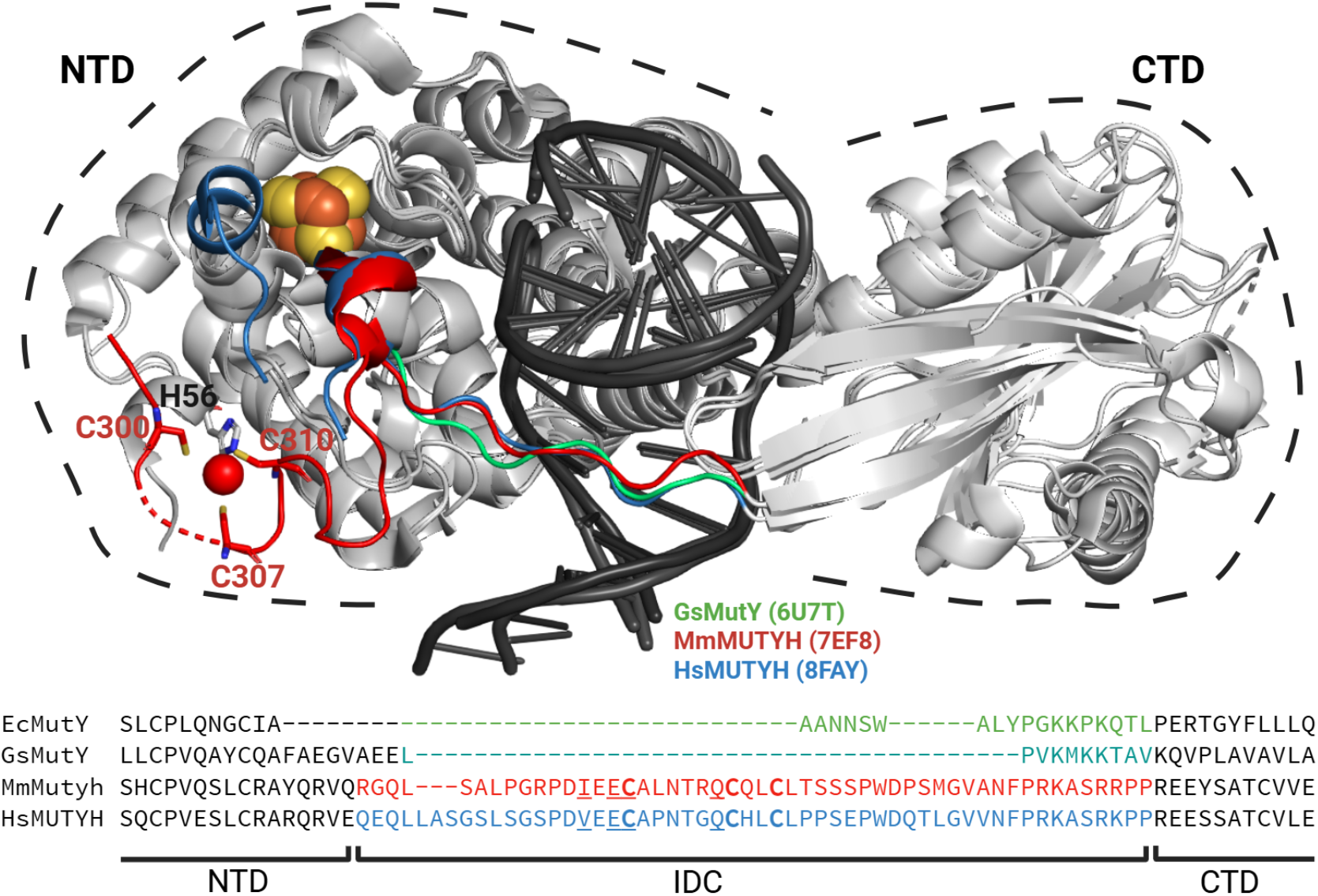
Differences in Interdomain Connector (IDC) of bacterial MutY and mammalian homologs Structure of GsMutY, MmMutyh, HsMUTYH are overlay with regions that are different highlighted in green, red and blue, respectively. The cancer associated variants studied herein are (mouse: NM_133250.2/human: NM_001128425.2): I297M/V329M, E299K/E331K, C300R/C332R, Q306H/Q338H, and Q306R/Q338R, the active site variant D207N/D236N, and the “founder” cancer-associated variants Y150C/Y179C and G365D/G396D. The pertinent residues in the IDC are underlined.

While the N- and C-terminal domains are spatially separated, their functions are connected through a hydrogen-bonding network comprised of residues in both domains. In bacterial MutY, residues in both domains are involved in OG recognition to ensure proper excision of only the inappropriately paired As by the N-terminal catalytic domain.[7] DNA binding via the [4Fe-4S] cluster domain of human MUTYH is also allosterically linked to the base excision site via an intricate H-bond network.[13] Amino acid residues may play multiple roles in enzyme function, such that a mutation of a single residue may have both local and long-range impacts. Indeed, a remarkable 85% of residues in proteins have been suggested to mediate long-range interactions, indicating that it is highly likely that a mutation may alter more than just the local structure and stability.[19] For example, a mutation may disrupt long-range contacts between different regions of the protein that cause an increase in unfolded protein entropy, thereby leading to destabilization of the folded protein.[20]

Herein, we selected several catalogued MUTYH variants in the IDC (Figure 2C) for functional testing to reveal how these amino acid changes affect various aspects of MUTYH-mediated repair. Specific mutations within the IDC that were chosen for this study include V329M, E331K, C332R, and Q338H/R/X. Q338H is common and generally referred to as a polymorphism, though some studies associated this variant with cancer.[21] The prevalence of the Q338H mutation is particularly high, up to 49%, in Japanese populations.[22] The observation of other variants at this Gln residue also suggest functional importance of this region. In contrast, the other selected

MUTYH variants are rare, and therefore designation as pathogenic or benign cannot be determined from genetics alone.[23,24] Variant frequency may also be distinct between populations of different ancestry. For example, there is a high frequency of Y179C and G396D in European populations, but not Asian populations.[25] Moreover, a study in the Italian population revealed a lower frequency of Y179C and G396D, and variants of uncertain significance (VUS) in 10% of participants, including V329M and Q338X.[26] The V329M mutation alters position 6 of the conserved <u>SG</u>XY<u>DV</u> motif for APE1 binding, where the last residue may be any non-polar aliphatic residue.[15] Furthermore, Val329 has been shown to be required for interaction with HUS1.[17] Upon inspection of the proposed HUS1 binding region in the MUTYH IDC, Glu331 is also conserved in several homologs, suggesting potential disruption to binding by the mutation to Lys in E331K.[27] C332R is a mutation to the first Cys of the <u>Cys</u>X_6_CysX_2_Cys coordinating Zn^2+^ ligands and would be anticipated to be detrimental to Zn coordination.[10]

Functional impacts of the selected IDC variants entailed a coordinated set of *in vitro* assays and analysis using a newly optimized cellular repair assay. The variants were first evaluated *in vitro* through metal analysis, adenine glycosylase assays, APE1 stimulation assays, and fluorescence polarization binding affinity measurements. Surprisingly, in most cases, the selected IDC cancer associated variants (CAVs) resembled the WT enzyme, suggesting that potential defects may be too subtle to detect via these assays or that these variations alter other aspects of the repair process. Alternatively, though associated with cancer, these variants may be functionally benign. We benchmarked the newly optimized cellular repair assay by evaluating OG:A repair using the two well-characterized MAP variants Y179C and G396D, and catalytically inactive CAV D236N. Surprisingly, reduced cellular repair was observed for many of the IDC CAVs, despite resembling the WT protein *in vitro*.

## Results and Discussion

### Purification and metal analysis of IDC Mutyh variants

The *in vitro* glycosylase and binding parameters for each of the selected human MUTYH variants in the IDC were evaluated using the corresponding mutation in the *Mus musculus* protein, Mutyh (NM_133250.2). The IDC variants in Mutyh are I297M, E299K, C300R, Q306R/H/X. Of note, Val329 in human MUTYH corresponds to an Ile residue in Mutyh, preserving an aliphatic and nonpolar residue at this position. The Mutyh variants were purified as previously reported for the WT enzyme.[14,28] Notably, we were unable to isolate sufficient quantities of Q306X Mutyh for enzyme assays, likely due to the instability of the protein caused by the truncation at this residue.

Inductively coupled plasma-mass spectrometry (ICP-MS) analysis of the variant proteins was performed to discern impacts on [4Fe-4S] cluster and Zn cofactor retention due to the amino acid variations. The wild-type enzyme was shown to harbor approximately 4 mol Fe and 1 mol Zn per mol enzyme (Table 1), similar to previous reports.[10] The percent cluster loading was measured as the fraction of the total protein that contained a [4Fe-4S] cluster using the concentrations based on UV-Vis absorbance at 410 nm and 280 nm. Notably, we commonly observe MutY/Mutyh to be sub-stoichiometrically loaded with [4Fe-4S] and Zn, likely due to incomplete cofactor insertion or loss during purification.[13,29–31] To reflect these potential variations in analysis of cofactor retention, we also considered the ratios of Fe/Zn/Abs280. Taken all together, the ICP-MS analyses suggest similar retention of cofactors in the series of variants as in the wild-type enzyme (Table 1).

**Table 1:** Metal cofactor loading adenine glycosylase activity and lesion affinity of IDC variants.

| Mutyh | % cluster loading<sup>a</sup> | Fe<sup>b</sup> | Zn<sup>b</sup> | Fe/Zn ratio | Active Fraction (%)<sup>c</sup> | $k_2(\text{min}^{-1})^d$ | $K_{1/2}(\text{nM})^e$ |
| --- | --- | --- | --- | --- | --- | --- | --- |
| Wild type | 75 | $4.0 \pm 0.2^f$ | $0.9 \pm 0.01^f$ | $4 \pm 0.23^f$ | $45 \pm 1\%$ | $1.1 \pm 0.1$ | $47 \pm 9$ |
| I297M | 87 | $3.3 \pm 0.1$ | $1.3 \pm 0.1$ | $2.5 \pm 0.2$ | $31 \pm 3$ | $1.2 \pm 0.1$ | $17 \pm 2$ |
| E299K | 73 | $3.1 \pm 0.1$ | $0.5 \pm 0.1$ | $6.2 \pm 0.1$ | $29 \pm 7$ | $1.1 \pm 0.1$ | $51 \pm 10$ |
| C300R | 78 | $2.9 \pm 0.3$ | $1.2 \pm 0.1$ | $2.4 \pm 0.3$ | $28 \pm 1$ | $1.5 \pm 0.1$ | $26 \pm 1$ |
| Q306R | 75 | $2.9 \pm 0.2$ | $1.0 \pm 0.1$ | $2.9 \pm 0.4$ | $31 \pm 5$ | $1.1 \pm 0.1$ | $59 \pm 12$ |
| Q306H | 72 | $3.0 \pm 0.2$ | $0.8 \pm 0.1$ | $3.8 \pm 0.5$ | $47 \pm 9$ | $1.1 \pm 0.1$ | $26 \pm 3$ |

<sup>a</sup>[4Fe-4S] cluster loading is based on UV absorbance concentrations evaluated at 410 nm (Fe-S cluster) versus 280 nm (protein).<sup>b</sup>Fe and Zn content is based on ICP-MS data relative to protein concentration measured by absorbance. Fe/Zn ratio is calculated using the following equations: 1) Ratio (R): $\text{Fe/Zn}$ 2) Propagated relative error: $(\Delta R/R)^2 = (\Delta \text{Fe} / \text{Fe})^2 + (\Delta \text{Zn} / \text{Zn})^2$ 3) Absolute error: $\Delta R = R \sqrt{(\Delta \text{Fe}/\text{Fe})^2 + (\Delta \text{Zn}/\text{Zn})^2}$ <sup>c</sup>Active fraction is based on amplitude of burst phase from MTO experiments (consult methods) <sup>d</sup>Rate constants $k_2$ were measured under STO conditions at 37 °C. <sup>e</sup>Apparent dissociation constants, reported as $K_{1/2}$ , of Mutyh and IDC variants were determined using the substrate analog 2'-FA in a fluorescence polarization assay.<sup>f</sup>As previously reported[10].

### Adenine glycosylase activity and binding affinity of IDC Mutyh variants

To delineate potential *in vitro* functional defects in the series of Mutyh variants, we performed detailed adenine glycosylase and fluorescence polarization assays to measure kinetic and binding parameters, respectively. The adenine glycosylase activity of Mutyh and variants was assessed using previously established assays.[28,32,33] Briefly, glycosylase assays entailed monitoring the extent of strand scission via denaturing PAGE at the apurinic (AP) site generated at the A paired opposite OG within a 30 base pair duplex. All the variant enzymes displayed biphasic kinetic behavior under multiple-turnover conditions (MTO), consisting of a “burst” phase followed by a slow, linear phase of product formation, similar to that observed with the WT enzyme (Figure S1); we therefore used the same kinetic scheme and approach for measuring the kinetic parameters (Scheme 1) as we have previously reported.[28,32]

Multiple turnover (MTO) experiments ([E] = 2 nM to 8 nM, and [DNA] = 20 nM) were used to determine active fraction based on the amplitude of the burst phase. Production curves were fit with the appropriate equations (see Methods and Supporting information) to extract burst amplitudes and calculate an active fraction relative to the total protein concentration. Interestingly, the active fraction of several variants (I297M, E299K, Q306R) was lower (∼ 20-30%) than the wild type. However, reduced activity does not appear to correlate with loss of the metal cofactors (Table 1). The reduced active fraction may be a result of misfolding and/or inability to engage the OG:A substrate as effectively as the WT enzyme. Under single turnover (STO) conditions with 100 nM active wild-type or variant Mutyh and 20 nM DNA, the rate of glycosidic bond cleavage, *k*_2_, was found to be remarkably similar for all of the CAVs to that of the wild-type enzyme (Table 1).

**Scheme 1.**
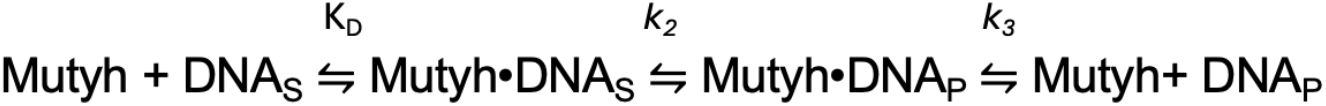
Minimal kinetics scheme for Mutyh.

Fluorescence polarization experiments were used to determine apparent dissociation constants (K_1/2_) of wild type and Mutyh variants with a 30 bp DNA duplex containing a non-cleavable 2’-deoxyadenosine analog, 2’-fluoro-2’-deoxyadenosine (FA) paired opposite OG. Overall, the variants bind to the substrate analog with a binding affinity of a similar order of magnitude as the wild-type enzyme (Table 1, Figure 3B), suggesting that these amino acid variations in the IDC do not substantially perturb lesion recognition by Mutyh.

**Figure 3:**
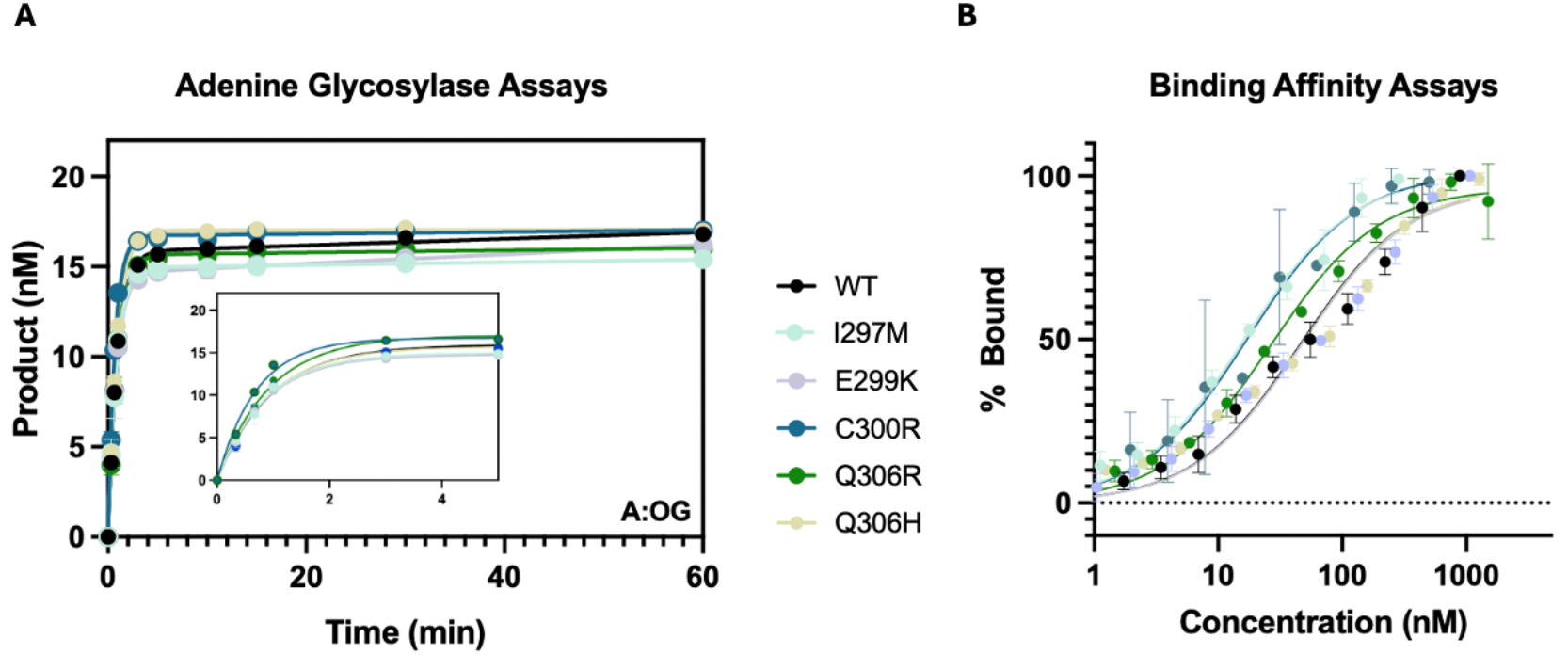
Adenine glycosylase activity and lesion affinity of WT and variant Mutyh A) Adenine glycosylase assay performed under single turnover conditions (STO) using 100 nM active wild-type or variant Mutyh and 20 nM OG:A-containing 30 bp duplex DNA (see sequence in Table S2) at 37 ºC. Data was fit to a single-exponential equation to determine rate constant *k*_2_ with a minimum of three trials; values are listed in Table 1. The inset expands the first 4 minutes to show the similarity of the variants to the WT B) Relative binding affinities were determined using the same 30 bp duplex containing the analog 2’-FA paired opposite OG using fluorescence polarization. Apparent dissociation constants (K_1/2_) were obtained by fitting with a one-site binding isotherm and are listed in Table 1.

The *in vitro* adenine glycosylase activity and lesion affinity of this set of Mutyh variants localized in the IDC are similar to the WT enzyme. Notably, this is reminiscent of what we observed with G396D MUTYH and the corresponding mutations in the bacterial and mouse proteins. In contrast, Y179C MUTYH and the corresponding bacterial and mouse variants, consistent exhibited a significantly compromised adenine glycosylase activity.[28,34] Both MAP variants exhibited compromised affinity for the OG:FA-containing duplex, which is also distinct from that observed with the IDC CAVs where no such binding defect was observed.[28] The impact of the Y179C and G396D variants on mismatch affinity and glycosylase activity is consistent with the structural information; Tyr179 plays a critical role in intercalation 5’ to the OG base to disrupt the bp and stabilize the flipped-out adenine, which Gly396 is located in a turn region that H-bonds to the OG phosphodiester backbone.[11,35,36] In contrast, the IDC is distal to the OG lesion recognition and adenine base excision site. Notably, the IDC is a reported locus for protein-protein interactions that may facilitate lesion recognition and hand-off to downstream repair enzymes.[15,27,37] Therefore potential impacts of variants on other aspects of the repair process may not be observed in glycosylase or binding assays, suggesting the need for analysis in different types of functional assays.

### AP endonuclease 1 (APE1) stimulation of Mutyh variant OG:AP-DNA product release

The location of the IDC Mutyh variants near or within the proposed APE1 binding site suggested potential impacts on APE1-stimulated substrate turnover.[27,38] In the absence of APE1, AP site product release rate constant *k*_3_for WT Mutyh is small (*k*_3_= 0.004 min^-1^) (Table 2). All IDC mutants exhibit similar rates of release of the AP site product in the absence of hAPE1. However, addition of 500 nM APE1 to the glycosylase reactions, enhances product release for all the IDC Mutyh variants (Figure S1).[28] In the case of WT Mutyh, APE1 enhances turnover over 10-fold (Table 2).[18] Product release was stimulated by the Mutyh variants to varying degrees indicating that the coordination with APE1 is retained, at least to some extent.

**Table 2:** Rate of product release, k_3_, of Mutyh in the absence and presence of 500 nM APE1^a^.

| Mutyh | $k_3$ without APE1 ($\text{min}^{-1}$) | $k_3$ with APE1 ($\text{min}^{-1}$) | Fold change |
| --- | --- | --- | --- |
| Wild-type | 0.004±0.003 | 0.04±0.01 | 10 |
| I297M | 0.003±0.002 | 0.03±0.001 | 10 |
| E299K | 0.002±0.001 | 0.020±0.003 | 10 |
| C300R | 0.0010±0.0007 | 0.020±0.001 | 20 |
| Q306H | 0.006±0.003 | 0.03±0.01 | 5 |
| Q306R | 0.0010±0.0008 | 0.020±0.004 | 20 |

<sup>a</sup> These experiments were conducted under multiple turnover conditions at 37 °C supplemented with 2 nM MgCl<sub>2</sub>. See methods for details of experimental conditions.

Of the mutants studied, V329M is the only one directly localized within the APE1 consensus binding motif.[15] The APE1 stimulation assay with the orthologous mutation in Mutyh, I297M, suggests that the APE1 interaction is minimally impacted, within the error of our measurement. The other CAVs studied do not reside directly within the proposed interaction motif but are anticipated to alter adjacent regions. The fold change of the rate of product release between the absence and presence of APE1 for the C300R and Q306R mutants are slightly higher than with the WT enzyme (Table 2). However, with these two variants, the product release rates, *k*_3_, in the absence of APE1 are smaller than WT (0.001 min^-1^), enhancing impact of APE1. In contrast, Q306H exhibits a slightly reduced enhancement of product release with APE1 compared to the WT enzyme, that suggests a less effective interaction with APE1.

In previous work with G365D Mutyh, we observed that the presence of 500 nM APE1 was less effective than WT in stimulating product formation (2-fold vs 10-fold). However, under these same conditions, rather than enhancing product formation, the presence of APE1 drastically inhibited product formation with Y150C Mutyh.[28] In this case, the amplitude of the burst phase was dramatically reduced suggesting that APE1 was competing with Y150C Mutyh for the OG:A DNA substrate. Indeed, both G365D and Y150C variants were inhibited under STO by the presence of APE1, suggesting that their reduced substrate affinity results in sensitivity of other DNA binding proteins. These results are in contrast with what we observe herein with the IDC variants where APE1 did not reduce the burst amplitudes and stimulated steady-state turnover.

### An assay for evaluating MUTYH-mediated repair in mammalian cells

To evaluate whether MUTYH IDC variants exhibit reduced activity in the context of a more complex cellular environment, an OG:A-specific repair assay in mammalian cells was developed (Figure 4). Stably-transfected MUTYH IDC variant mammalian cell lines were prepared and used to evaluate repair of an OG:A mismatch localized within a GFP plasmid reporter. Recently, we reported an improved method for generation of the OG:A containing lesion plasmid and showed that this could be used to define structure-activity relationships of MUTYH repair of X:A mismatches, where X = OG and synthetic OG analogs.[39] The improved plasmid reporter design has also enabled higher efficiency transfection, increased cell counts to measure repair, and a decrease in plasmid demand. Using this newly optimized assay, the presence or absence of GFP fluorescence from variant or WT expressing cells was monitored using flow cytometry (Figure 4). All transfected cells express RFP and only those that repair the OG:A lesion also express GFP. Percent repair of OG:A in each cell line was normalized to the RFP+/GFP+ positive control.

**Figure 4:**
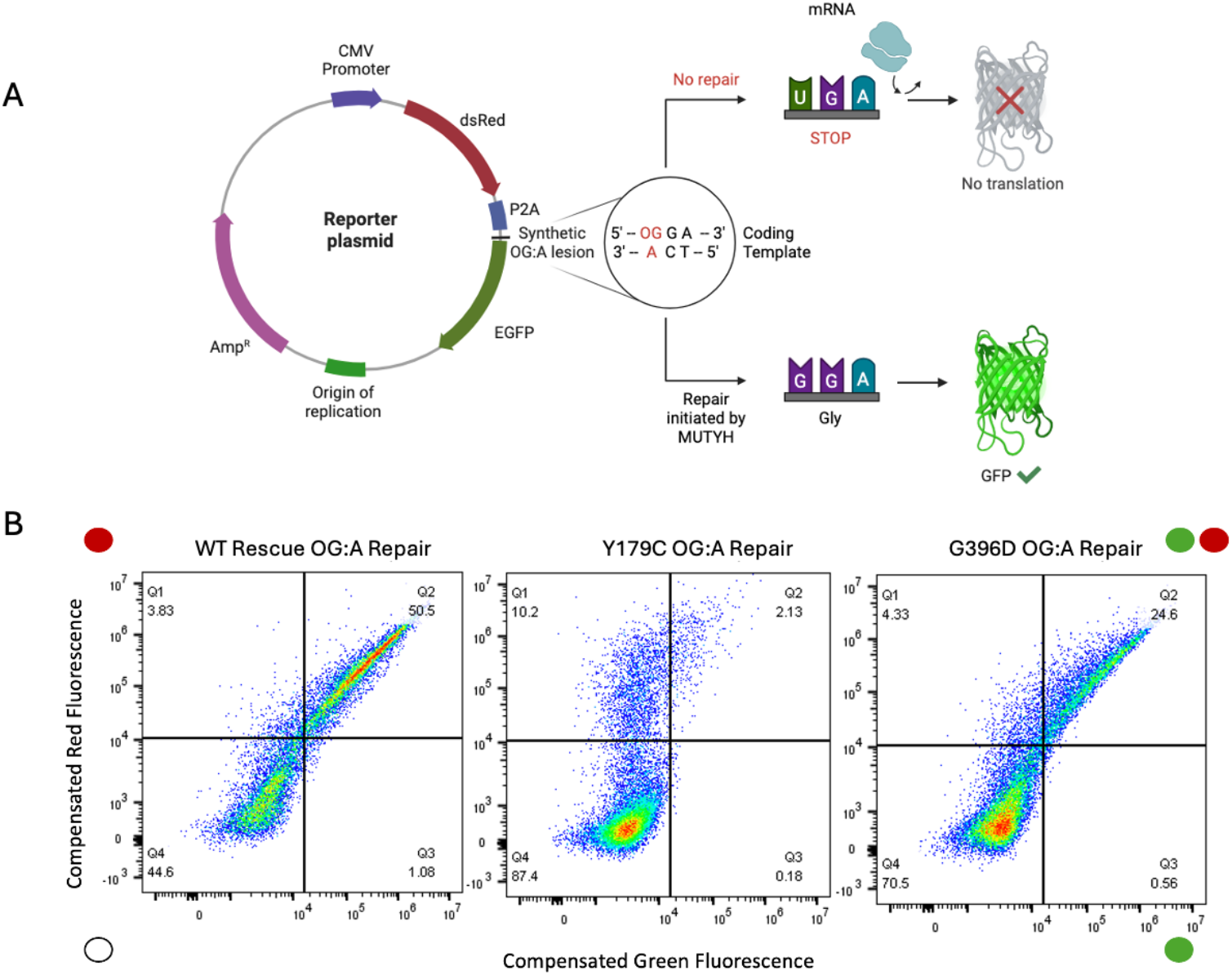
OG:A Repair Plasmid and cell assay design for analysis of MUTYH-mediated repair in cells. A) The reporter plasmid used in the mammalian cell assay contains the coding regions for two fluorescent proteins: dsRed and GFP. The dsRed serves as a transfection control, whereas GFP expression is used to measure OG:A repair. If the synthetically inserted OG:A lesion is not repaired, the transcript encodes a stop codon and no GFP protein is produced. If the OG:A base pair is repaired back to G:C, the full-length protein is produced. B) Representative flow cytometry plots for OG:A repair in the WT Rescue, Y179C, and G396D cell lines, represented as compensated red (Y axis) vs green (X axis) fluorescence. The percentage in each quadrant represents the percentage of cells within that population, with Q1 being dsRed+, Q2 being dsRed+GFP+, Q3 being dsRed-GFP-(un-transfected cells), and Q4 being GFP+ cells.

The assay design was validated by stably transfecting the WT *MUTYH* gene into CRISPR-Cas9 generated knockout (KO) *MUTYH*^*-/-*^ human embryonic kidney (HEK)293FT cells to rescue OG:A repair. The HEK293FT cell lines possess an FRT site for single stable integration of the *MUTYH* gene. For all proteins studied, three individual colonies were each expanded to make three cell lines to assess repair in the mammalian cell assay. The human MUTYH type 2 (beta 3) 521-nuclear isoform was utilized to study the wild-type protein.[23] For consistency with previous studies, amino acid numbering reflects the 549 isoform. Introduction of the WT or variant MUTYH gene in the cell lines was confirmed using reverse-transcriptase polymerase chain reaction (RT-PCR) and Western blotting (Figure S2). Repair for each of the three clonal cell lines was assessed at least three times, and the percent repair reported is an average between all trials. Data is calculated as a ratio of the cells that express GFP and dsRed, divided by the total number of cells, and is represented as a percentage of repair of OG:A lesions ([dsRed+GFP+]/dsRed+*100). Remarkably, these WT rescue cells displayed levels of repair indistinguishable from the unmodified WT HEK293 cells (Figure 5A, B, S3). When quantified, the extent of OG:A repair in the WT rescue cells approached 100% without statistically significant differences from unmodified WT HEK293 cells (Figure 5A, B, S3).

**Figure 5:**
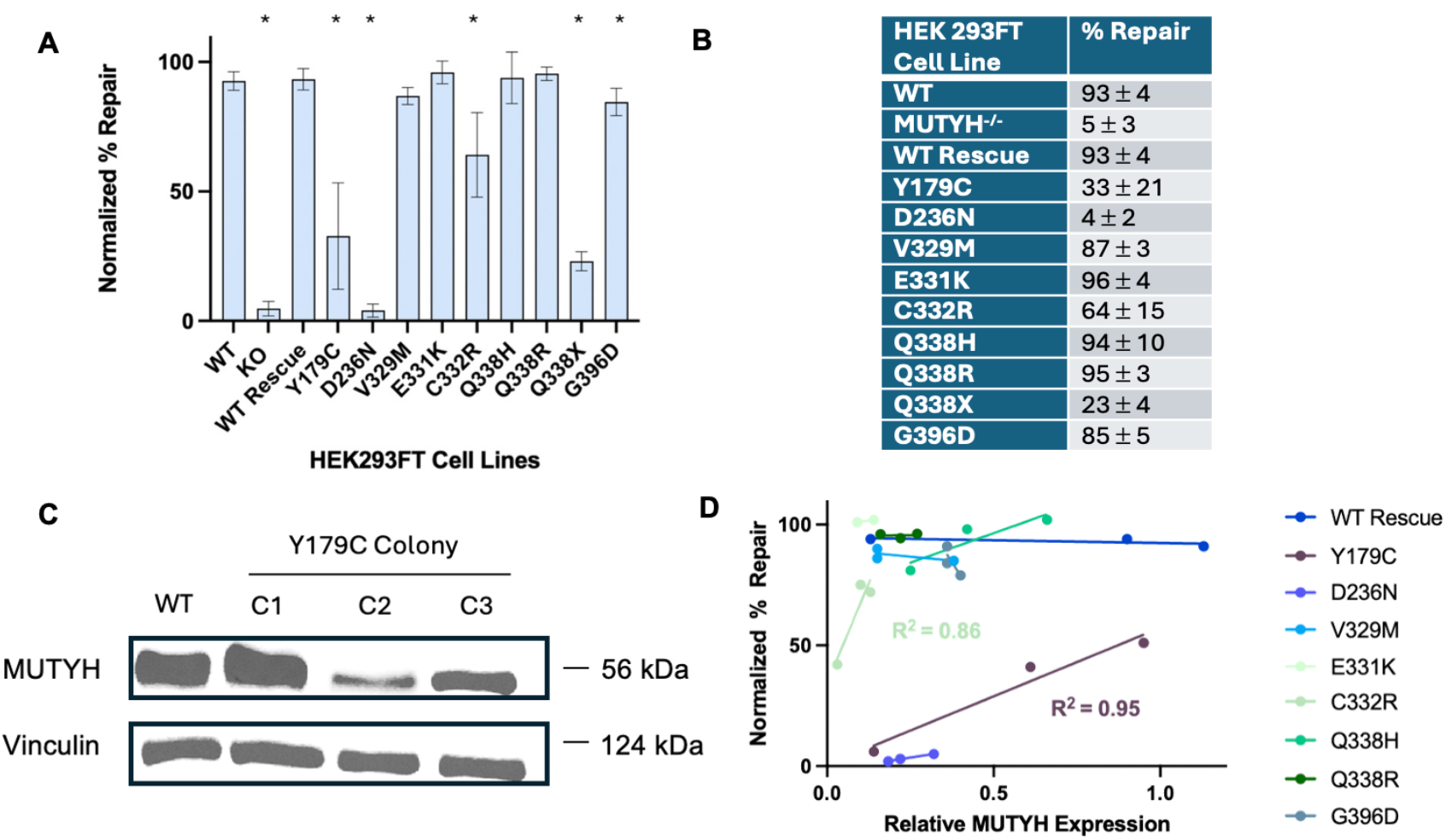
Normalized % OG:A Repair for MUTYH variants studied in this work. A) Data from each mutant was compiled and statistical significance was determined *via* student t-test. Each bar is an average of at least 9 trials. An asterisk indicates p < 0.05 *via* student t-test when compared to WT HEK293 cells (not indicated for KO cells). B) Numerical average % repair values plotted in (A) from each cell line (± standard deviation from average). C) Representative western blot image of three Y179C mutant clonal cell lines were generated simultaneously, but protein expression for these variants varied between clones despite recombination into the same genomic locus. Quantification of MUTYH protein is reported as a ratio of MUTYH to the housekeeping vinculin protein. All mutant cell lines were also quantified against the endogenous WT MUTYH protein and represented as protein expression relative to the endogenous WT protein. D) Correlation between relative MUTYH expression and normalized % OG:A repair. A simple linear regression analysis was plotted, and significant R^2^ values are reported. There is a positive correlation between protein expression and A:OG repair capacity by variants with a modest defect in repair capability, such as C332R and Y179C.

### Repair of OG:A mismatches mediated by MUTYH variants in human cells

To benchmark the OG:A repair levels in the cellular assay for the IDC CAVs, we evaluated the OG:A repair of cells expressing the MAP variants Y179C and G396D, and the catalytically inactive CAV D236N. The Y179C and G396D variants were the first to be identified in patients exhibiting colonic polyposis who did not possess mutations in *APC* and have since been robustly documented based on genetics as pathogenic MUTYH variants.[1] The human D236N variant, and its bacterial and mouse homologs, have been shown to be completely devoid of activity with OG:A substrates,[11,40] consistent with the critical role of this Asp as the nucleophile in the adenine glycosylase mechanism.[11,41] Using the newly optimized mammalian cell repair assay, the Y179C mutation resulted in significantly decreased OG:A repair (33 ± 21%) in all replicate cell lines compared to WT HEK293 cells (93± 4 %) (Figure 5A, B, S3). In contrast, the G396D mutant showed a more modest but statistically significant decrease in OG:A repair (84 ±5 %) compared to WT cells. Indeed, when averaged between all trials for each mutant, both mutants displayed statistically significant decreased OG:A repair from WT HEK293 cells (Figure 5A, B). The D236N variant cell lines exhibited even lower levels of OG:A repair (4 ± 2%) than Y179C cell lines, similar to repair levels observed in *MUTYH*^-/-^ HEK293 cells (5 ± 3%); this result confirms the expectation that a catalytically inactive enzyme is unable to mediate OG:A repair. These results illustrate the ability to observe a range of repair activities and detect defective repair of MUTYH variants in cells.

We next prepared the IDC variants expressing HEK cell lines and evaluated the ability of the IDC variants to mediate repair of the OG:A lesion within the reporter plasmid. Repair for each of the three clonal cell lines per variant was assessed at least three times, and the percent repair reported is an average between all trials, as illustrated with the other variants and WT rescue discussed above. Of the novel CAVs studied, half showed statistically significant differences in percent OG:A repair from WT HEKs, while the other half did not (Figure 5A, B, S3). The mutations V329M, C332R, and Q338X showed decreased OG:A repair of 87 ± 3, 64 ± 15, 23 ± 4%, respectively, compared to the WT mediated repair levels of 93 ± 4%. The Q338X nonsense mutation truncates MUTYH within the CTD OG recognition domain; therefore, we anticipated that the OG:A repair mediated in this cell line would be like that in MUTYH KO cells. Surprisingly, the observed repair levels mediated by Q338X (23 ± 4%) are above the levels observed in the KO cells (5 ± 3%). Mutations E331K, Q388H, and Q338R showed no significant difference from wild-type levels of repair, suggesting that these variations may not compromise MUTYH-mediated OG:A repair.

### MUTYH variant expression versus repair

In some cases, we observed differences in OG:A repair in three different cell lines expressing the same mutation (Figure S4). To delineate if this is a result of different levels of MUTYH protein within each cell line, we measured protein expression using quantitative Western blotting (Figures 5C, S2, S4). Specifically, protein expression for each cell line was analyzed in triplicate and normalized to the vinculin loading control. Average expression in each generated cell line was then compared to the endogenous WT MUTYH expression in HEK293FT cells.

We observed a correlation between cell lines relative expression level and OG:A repair levels for Y179C and C332R. Specifically, for Y179C Colony 2, less MUTYH protein is expressed than in colonies 1 and 3 (Figure S4, 5C, 5D), and less repair for colony 2 (6 ± 2%) is observed than for colonies 1 and 3 (51± 1 and 41±1 %, respectively). For C332R, there is also a positive correlation between expression and repair capacity, in which we observed the lowest average OG:A repair percentage in colony 2 (about 42 ± 1% as opposed to 72 ± 1% and 75 ± 8% for colonies 1 and 3, respectively). This supports the expectation that there is direct correlation between DNA glycosylase protein expression and activity that is variant specific. Notably, the correlation of repair capacity and MUTYH expression is only observed with those that show a major defect in repair capabilities. For example, relative expression and repair levels for G396D are similar across the different colonies (percent repair∼ 85%). In fact, the vast majority of MUTYH mutant cell lines, as well as the WT rescue, exhibit near 100% repair in all triplicate cell lines, including those expressing the lowest levels of protein. Comparing the cell lines with the lowest levels of expression more clearly magnifies the defective repair activity of C332R and Y179C.

We anticipated that repair in cells of variants at crucial catalytic residues, like D236N, would not be strongly influenced by protein expression levels. With the D236N cell lines, the levels of OG:A repair remained low (∼4%) across the expression range. In contrast, variants with more moderate defects (Y179C and C332R) show expression-dependent increases in repair activity well above baseline, indicating that their catalytic defects can be compensated for by increased expression. Taken together, these results suggest that some compromised MUTYH variants may display repair levels that are highly dependent on the level of expression, while others, like D236N, may be unable to mediate significant repair even when expressed at high levels. Mildly dysfunctional variants may be competent enough to mediate full repair even at low levels of expression.

Recently, a study using our OG:A repair strategy was leveraged to perform a high-throughput saturation mutagenesis study of MUTYH.[42] The overall results were consistent with what we observe in examining OG:A repair of cell lines of individual variants, but there were some differences. Notably, the high-throughput approach does not allow for a correlation to be made between repair and expression. For example, the pathogenic variant, Y179C was reported to exhibit no significant repair in their analysis, while our results show that this variant has reduced levels of repair relative to WT but that are above background. In addition, the hampered repair activity increases in clones with higher levels of Y179C expression. It may be anticipated that Y179C expression may be sensitive to cell type and conditions, thereby impacting overall repair. The high throughput analysis also reported mid-range repair levels (43%) for G396D, while we observe values closer to WT (about 85%). This may be due in part to the method of the analysis (FACS of mixtures of variants versus analysis of a single variants clonal cell lines) and differences in dynamic range and normalization methods. Another recent study[43] used CRISPR-Cas9 base editing methods to make several variants within the MUTYH gene in HEK293 cells and also used our OG:A reporter strategy, though none of the selected variants were identical to those we studied herein.[23,39,43] Of note, this approach would be anticipated to provide expression levels that match more closely those of endogenous MUTYH. We believe that it is important to consider potential differences in expression, which may be variant specific and differ among individuals and populations when drawing conclusions on variants based on cell repair assays.[44–47]

### Structure Activity Correlations for IDC Variants

We uncovered differences among the IDC variants in the cellular repair assay, despite their similarities in the *in vitro* glycosylase activity, binding affinities, and APE1-mediated product release rates to the WT enzyme. The cellular repair assays with V329M, C332R, and Q338X showed significant reduction in OG:A repair. These mutations alter regions of the protein known to be important for protein-protein interactions, metal coordination, and substrate recognition, respectively. Curiously, the truncated Gln338X variant exhibited higher levels of OG:A repair than the *MUTYH* KO cells. This suggests that a small amount of truncated MUTYH protein may be present to mediate above background levels of OG:A repair. We were unsuccessful in overexpressing and purifying the corresponding Mutyh variant, suggesting that the truncating mutation destabilizes protein stability. However, our analysis of expression versus repair (Figure 5 C, D) suggests that only low levels of WT and some MUTYH variants are needed to mediate full repair. This mutation truncates MUTYH within its C-terminal OG recognition domain, and this truncated enzyme may retain activity. In fact, in previous studies we demonstrated a similarly truncated bacterial protein, MutYΔCTD was catalytically active despite lacking the C-terminal OG recognition domain.[34] Alternatively, the small amount of repair observed with Q338X may be due to read-through of the stop codon, though we were unable to detect full-length protein by Western Blot analysis [48]. Nevertheless, the extent of OG:A repair mediated by this variant is near that of the MUTYH KO. This mutant is annotated as pathogenic by ClinVar, and results described herein support that hypothesis.[6]

The C332R variant exhibited significantly reduced activity in the cellular assay (64 ± 15 %), approaching that of the known cancer variant Y179C (33 ± 21 %). In contrast to Y179C, which exhibited significantly reduced activity in vitro, the corresponding mutation in the mouse Mutyh (C300R) exhibited adenine glycosylase activity similar to the WT enzyme.[1,3,28] C300R Mutyh also retained full loading of Zn^2+^, which was especially surprising since Cys300/332 corresponds to one of the Zn^2+^ coordinating ligands. In previous work, mutations of Cys300 to Ser was shown to not alter metal loading, suggesting that this particular Cys residue is more weakly coordinating than the other Cys ligands within the IDC. However, we anticipated that replacing the Cys residue with a large positively charged residue in the vicinity of the positively charged Zn ion would be unfavorable for Zn coordination. The ability to accommodate the Cys-to-Arg mutation may suggest that the Zn coordination site is flexible and may be related to the highly solvent exposed nature of the IDC. Notably, in the reported structures, the IDC is highly disordered and therefore the impact of the Arg mutation is not readily predicted based on structures alone.[8,12] The more dramatic impact on MUTYH-mediated repair function in cells suggests that the impact on metal coordination and/or folding of the Zinc Linchpin may impact lesion engagement and repair more dramatically in cells. Indeed, in studies with the bacterial MutYs, we have previously observed that amino acid alterations in regions involved in lesion detection, such as within the FSH loop of the CTD, are more dramatically impacted in a cellular context where locating the lesion is more challenging due to its rarity and competition with other cellular proteins and processes.[49,50]

C332R and V329M may also alter steps downstream of base excision repair, particularly the recruitment and handoff to APE1. The *in vitro* adenine glycosylase activity of the V329M variant, like C332R, resembled that of the wild-type protein, but exhibited OG:A repair activity in cells more closely resembling that of the G396D variant. The IDC has been identified as a docking site for many different protein-protein interactions, so it is likely one or more of these would be affected by a variant in this region despite maintaining wild-type adenine glycosylase activity. Both mutations may affect product handoff from MUTYH to APE1 which would be required in our assay to observe repair. As the readout of this assay is dependent on full conversion from OG:A back to a G:C pair, lack of GFP fluorescence does not necessarily reveal the aspect of overall repair affected by these variant, but taken together with the in vitro analysis implicates the APE1 hand-off step. Since these mutations reside within or adjacent to known protein-protein interfaces, additional pull-down or crosslinking experiments would aid in identifying impaired interactions. Furthermore, these mutations may affect roles of MUTYH that have yet to be characterized, such as its interactions with HUS1.[51] Thus, additional studies will be required to identify which interactions may be perturbed by amino acid variants.

Other mutants studied (E331K, Q388H, and Q338R) displayed only slightly decreased cellular OG:A repair relative to cells expressing WT MUTYH. These variants also resemble the wild-type enzyme in vitro. Interestingly, the Q338H mutation appears to resemble wild-type in this report in contrast to our earlier studies.[23] Previously, our laboratory reported lower OG:A repair of Q338H compared to wild-type, but greater than Y179C in Mouse Embryonic Fibroblasts (MEFs).[23] The difference with our results herein may be due to plasmid design and cell types. For example, MEF cells are more sensitive to standard culture conditions than human embryonic fibroblast cells and prematurely enter cellular senescence due to increased oxidative stress.[52] Along this line, a previous study of *Mutyh*^-/-^ MEFs expressing Q338H were shown to be hypersensitive to treatment with KBrO_3_and exhibited an arrested progression through S-phase, along with increased levels of OG suggesting a repair defect.[24] This prompts an intriguing hypothesis that Q338H, along with other variants, may display a different phenotype in high *versus* basal reactive oxygen species (ROS) conditions. High ROS levels are associated with inflammation and cancer, and these may exacerbate defects in OG:A repair of MUTYH variants.[53–56] Additional studies will be needed to determine how variants function under a range of ROS conditions and whether that may contribute to a polyposis phenotype. In addition, mild defects in OG:A repair may only be problematic later in life. Reduced DNA repair capacity has been associated with aging, and when combined with accumulated DNA damage may explain how cancer phenotypes develop late in life.[57] Additional studies will be required to determine how cellular conditions, mutations in other genes, and environmental stressors impact OG:A repair and impact disease progression in a given individual.

## Conclusions

The study herein illustrates the power of combining in vitro and cellular assays to reveal functional information and clinical impact of *MUTYH* variants. A summary of the functional data and insight based on structural information for the IDC CAVs along with Y179C, G396D and D236N MUTYH is shown in Figure 6. Taking the structural, functional, and cellular assay data together suggests dysfunction by all metrics (red) of G338X, Y179C and D236N (except for lesion affinity), while E331K and Q338H/R appear like WT (green) by these same metrics. The other variants are mildly compromised (yellow/salmon) in a select categories, though notably similar to what has been found with the well-studied MAP variant G396D. This detailed analysis with a variety of assays provides additional confidence in categorizing these variants not just as pathogenic or benign, but also with resolution for having mild or mid-range impacts. Knowledge of variant dysfunction and its magnitude can be added to the information provided to clinicians and patients in assessing cancer risk. Cellular OG:A reporter studies may be particularly useful in providing an indication of compromised variants; subsequent in-depth biochemical studies may then be performed to identify the specific enzymatic function(s) that are altered by the MUTYH variant. Information from a variety of assays not only provides confidence in clinical designation but also detailed insights into the many discrete steps involved in MUTYH-mediated repair.

**Figure 6:**
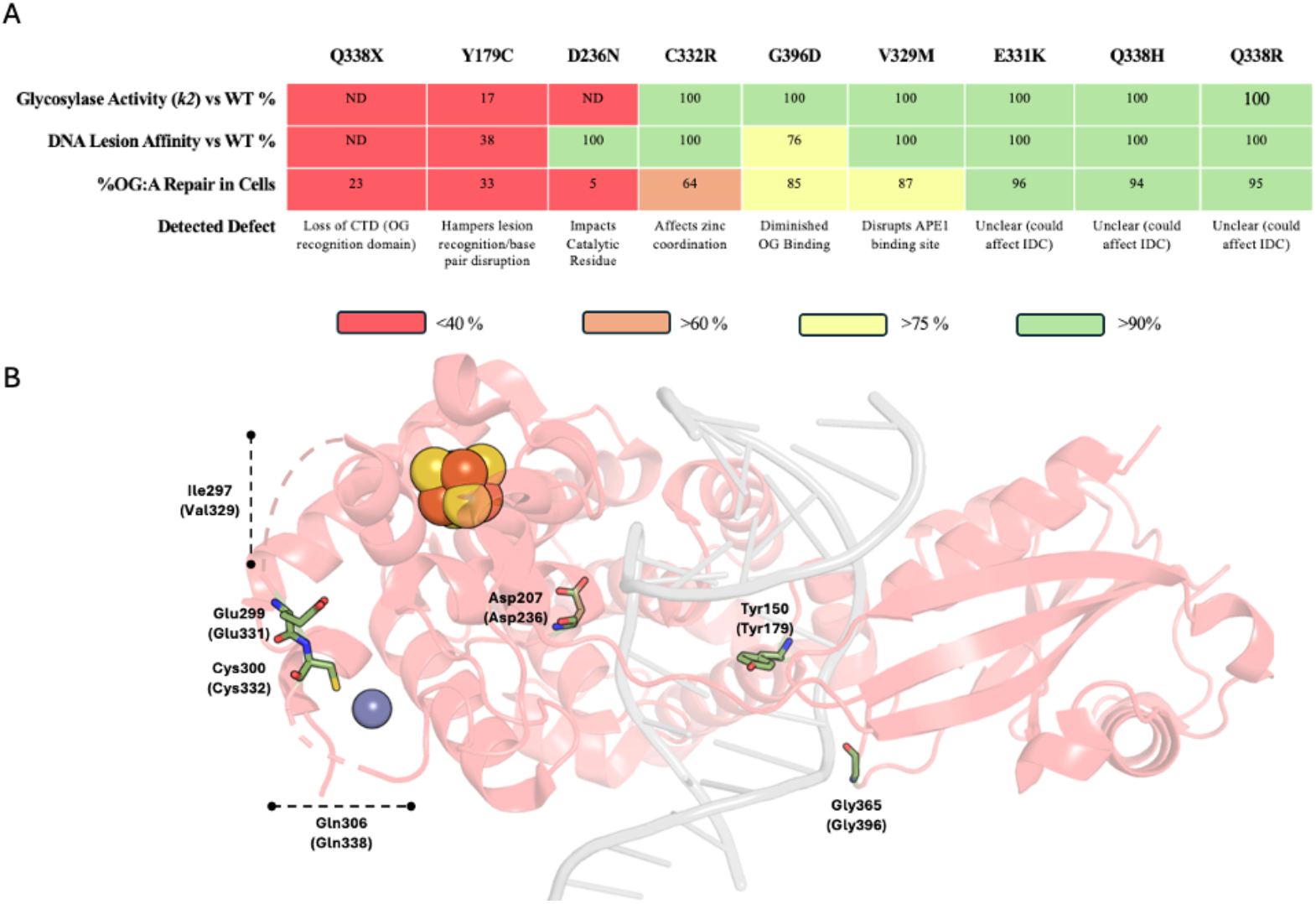
Functional Consequences and Structural Context of MUTYH Cancer-Associcated Variants. A)Summary of glycosylase activity, DNA binding affinity, and cellular repair capacity for all MUTYH variants examined. Major defects are indicated in red, moderate defects in orange, minor defects in yellow, and absence of detectable defects are green. Data for Y150C, G265D and D236N Mutyh was previously reported^[28][54]^ B)Crystal structure of Mutyh bound to DNA, highlighting residues corresponding to the cancer-associated variants analyzed herein. Mouse residue numbering (NM_133250.2) is shown above, with the corresponding human numbering (NM_001128425.2) indicated in parentheses.

### Experimental Section

Detailed experimental methods are provided in the Supporting Information

## Acknowledgements

This work was supported by the National Cancer Institute of the National Institutes of Health (CA067985 to S.S.D.) CHTA was supported in part by a postdoctoral fellowship from UC-MEXUS. M.M. and C.K. were supported by a National Institutes of Environmental Health Sciences (NIEHS)-funded predoctoral fellowships (T32 ES007059). C.K was also supported in part by a Floyd and Mary Schwall Dissertation Year Fellowship in Medical Research from UC Davis Graduate Studies. S.G.C. was supported in part by an ARCS Foundation Fellowship. G.P.W. was supported by a UC Davis Chemical Biology Program supported by an NIGMS Institutional Training Grant (T32 GM136597). We thank Dr. Nicole Nuñez and Dr. Alan Raetz for making the C300R and Q306H *pET28a* plasmids, the Heyer lab (UCD) for help with CsCl purification, and Prof. Jeff Miller (UCLA) for providing the *Mutyh* gene within the pQE30 plasmid.

## Supporting Information

Additional supporting information can be found online in the Supporting Information section: **Methods**; **Supporting Fig. S1**: Stimulation of MUTYH variants by APE1; **Supporting Fig. S2**:. Western Blot Confirmation of Generated Cell Lines; **Supporting Fig. S3**. Normalized OG:A % Repair per Colony; **Supporting Fig. S4**. Relative MUTYH expression in selected clones of generated cell lines; **Supporting Table S1:** Primers used to create mutants used in this study; **Supporting Table S2**: Oligonucleotide sequences; **Supporting Table S3**: Primers used for Cell Line Validation

## Table of Contents Graphic and Text

Functional analysis of cancer-associated variants of the DNA repair enzyme MUTYH in the interdomain connector between the 8-oxoguanine (OG) recognition and base excision domains reveals discordance between *in vitro* assays and OG:A repair in cells. This disconnect highlights use of complementary biochemical and cellular assays to accurately classify variant dysfunction.

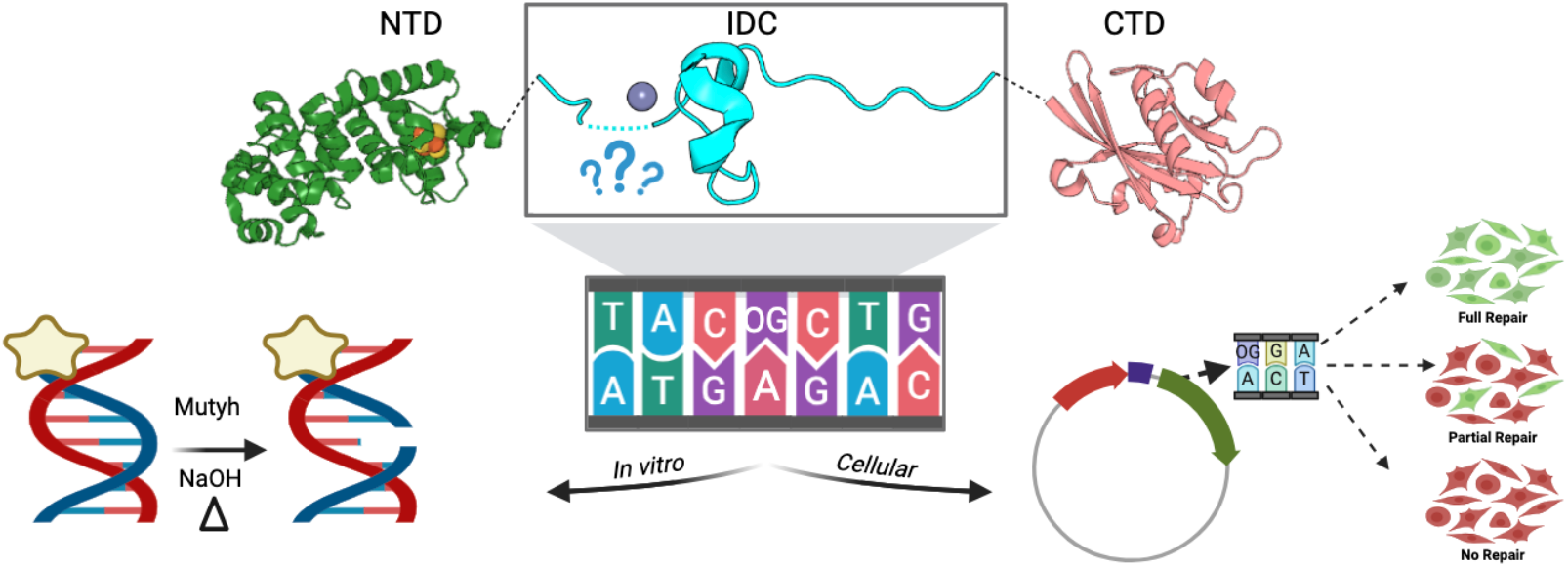

