## Supplementary Information for "MUTYH cancer-associated variants within the interdomain connector differentially impact glycosylase activity and cellular DNA repair"

__Table of Contents__ __Pages__

**Methods** **2-7**

**Supporting Fig. S1**: Stimulation of MUTYH variants by APE1 **7**

**Supporting Fig. S2**: Western Blot Confirmation of Generated Cell Lines **8**

**Supporting Fig. S3**. Normalized OG:A % Repair per Colony **8**

**Supporting Fig. S4.** Relative MUTYH expression in clones of generated cell lines **9**

**Supporting Table S1:** Primers used to create mutants used in this study **9**

**Supporting Table S2**: Oligonucleotide sequences **10**

**Supporting Table S3**: Primers used for Cell Line Validation **10**

**References** **11**

### Methods

*Site directed mutagenesis*

*In vitro* experiments were performed using the purified *Mus musculus* MutY homolog protein, Mutyh. The *Mutyh* gene within a pQE30 vector lacking the first 14 amino acids and containing a 6x-His tag with a thrombin cleavage site[1] was cloned into a *pET28a* vector for overexpression and purification.[2] Mutations were created through site directed mutagenesis using splicing by overlap extension (SOE) PCR or QuikChange (NEB). The primers used to create each mutant are listed in Table S1. Reactions for each fragment were created with either the FlankingFw plus the reverse (Rv) mutation primer or the FlankingRv plus the forward (Fw) mutation primer. Briefly, these reactions were carried out with 1x Phusion reaction buffer, 0.2 mM dNTPs, 0.5 µM of each primer, and 20-40 ng of the WT Mutyh plasmid. The reactions were incubated at 98 ºC x 30 sec; 35 cycles of 98 ºC x 10 sec, 72 ºC x 30 sec, 72 ºC x 30 sec; a final incubation at 72 ºC x 5 min followed by a 4 ºC hold. Expected fragments were 843 bps for MutyhFlankingFw + Rv mutation primers and 680 bps for MutyhFlankingRv + Fw mutation primers. Fragments were run on a 1% agarose gel and gel purified using the Qiagen Gel Extraction kit according to the manufacturer's protocol. After gel purification, the gene was assembled using the same PCR conditions and the pair of gene fragments for each mutation. The assembled gene was digested with *Not*I and *Sal*I to create sticky ends required for ligation into the *Not*I and *Sal*I double digested *pET28a* plasmid. Ligation was carried out at 16 ºC for 16 h. The ligation product was transformed into chemically competent DH5$\alpha$ cells and single colonies were selected and grown up for validation by Sanger sequencing.

*Protein overexpression and purification*

Protein purification of Mutyh was carried out based on previously published protocols.[3] Briefly, the Mutyh carrying *pET28a* plasmid (Kan$^R$) was transformed into BL21 DE3 cells containing the *pRKISC* (Tet$^R$) Fe-S cluster assembly complex plasmid. A single colony was selected to inoculate an overnight Luria broth (LB) culture supplemented with 15 µg/ml tetracycline and 34 µg/ml kanamycin and grown at 37 ºC and 220 rpm. 48 ml of the overnight culture was added to each 2 L of LB supplemented with the same antibiotics and grown at 37 ºC and 180 rpm. Once the $OD_{600nm}$ reached ~0.8, overexpression was initiated by the addition of 1 mM IPTG and 0.1 g ferrous ammonium sulfate. The temperature was reduced to 30 ºC for overexpression and harvested after 6 h by centrifugation at 5,000 rpm for 20 min at 4 ºC. Pellets were resuspended in resuspension buffer, comprised of 20 mM sodium phosphate buffer pH 7.5 and 10% glycerol. A crushed protease inhibitor tablet was added to each bacterial pellet prior to storage at -80 ºC until purification.

The cell pellet was thawed at 37 ºC at 150 rpm for 30-45 min, lysed by sonication, and centrifuged at 10,000 rpm for 15 min at 4 ºC. The supernatant was decanted and 20 mM imidazole and 1 M NaCl were added. The solution was incubated with pre-rinsed $Ni^{2+}$-NTA resin (Qiagen) for 1 h with gentle rotation on a carousel at 4 ºC. The solution was poured into a fresh PD10 column. After the unbound proteins were allowed to flow through the column by gravity, the protein bound resin was washed with 20 ml of wash buffer, comprised of 20 mM sodium phosphate buffer pH 7.5, 10% glycerol, 1 M NaCl, and 20 mM imidazole. After washing, the protein was eluted twice with 3 ml of elution buffer (20 mM sodium phosphate buffer pH 7.5, 10% glycerol, 200 mM NaCl, and 500 mM imidazole). The combined eluant was diluted to 30-40 ml with Heparin A buffer (20 mM sodium phosphate buffer pH 7.6, 1 mM EDTA, 5% glycerol) and incubated overnight at 4 ºC with 2 U thrombin (Novagen) per mg of Mutyh to cleave the 6x-His tag.

After overnight incubation, final concentrations of 1 mM PMSF and 1 M NaCl were added to the solution and incubated for 1 h. The solution was simultaneously concentrated to 5 ml using an Amicon stirred cell concentrator loaded with a 10,000 MWCO filter. After concentration, the protein was batch bound to clean $Ni^{2+}$-NTA resin and incubated at 4 ºC for 1 h before pouring over a fresh PD10 column and collecting the flowthrough. The resin was washed with 2 ml wash buffer, and the flowthrough was collected. Both flowthroughs were combined and diluted to 40 ml using Heparin A buffer and filtered through a 0.2 μm filter. The protein was further purified on a 1 ml Hi-Trap Heparin column (GE Healthcare) on an AKTApurifier FPLC using a linear gradient from 0-100% Heparin B (20 mM sodium phosphate buffer pH 7.6, 1 mM EDTA, 5% glycerol, 1 M NaCl) over 35 column volumes. Peak fractions corresponding to the full-length protein were collected and concentrated using an Amicon stirred cell concentrator with a 10,000 MWCO filter. The final protein volume was doubled with 50% glycerol, aliquoted into single use aliquots for *in vitro* studies and metal analysis, and stored at -80 ºC. Purity of each protein was assessed by a 12% SDS-PAGE stained with Sypro Orange according to the manufacturer's protocol. The concentration of total protein was determined by UV-vis using $\varepsilon_{280}$ = 86,120 $cm^{-1}$ $M^{-1}$ and $\varepsilon_{410}$ = 17,000 $cm^{-1}$ $M^{-1}$.

*Preparation of oligonucleotide substrates*
A 30-bp duplex (Duplex I) was used for i*n vitro* glycosylase assays and a 29-nucleotide OG-containing oligonucleotide (GFP OG) was used to create the OG:A mispair in the GFP reporter (sequences shown in Table S2). The A containing complementary strand was purchased from Sigma Aldrich. The OG and FA containing strands were synthesized by the University of Utah DNA and Peptide Synthesis Core Facility or IDT. For fluorescence polarization experiments, the FA containing complementary strand also contained a 5' label of a 6'-carboxyfluoresceine (6'FAM). The OG and FA containing strands were cleaved from the solid support column by incubation with $NH_4OH$ and ß-mercaptoethanol prior to HPLC purification. All oligonucleotides were purified on a Shimadzu HPLC with a Dionex 100 ion exchange column using a linear gradient of 0-2 mM ammonium acetate in 10% acetonitrile. Peak fractions were individually lyophilized and desalted with a SEP-PAK C18 column. Integrity of the DNA oligonucleotides was determined using MALDI-MS prior to use in experiments. Oligonucleotides were stored dried. Concentrations of solutions using calculated $\varepsilon_{260}$ values. Prior to use, the GFP OG oligonucleotide was phosphorylated using PNK kinase (New England Biolabs) according to the manufacturer's protocol.

*Adenine glycosylase assays*
*In vitro* glycosylase assays were performed as previously described.[3] Briefly, the A containing strand was radiolabeled with g-$^{32}$P using T4 kinase. After labeling, the strand was dried and annealed to the OG containing strand such that 5% of the DNA was radiolabeled. For glycosylase assays, the reaction mixture was prepared to final concentrations of 20 nM DNA in 20 mM Tris-HCl pH 7.6, 1 mM EDTA, 0.1 mg/ml BSA, and 30 mM NaCl and incubated at 37 ºC. Under multiple turnover conditions, the reaction was initiated by addition of 2-8 nM *Mutyh*. Under single turnover conditions, the reaction was initiated by addition of 100 nM active enzyme. An 8 μl aliquot of the reaction was quenched with 2 μl 1 M NaOH at time points ranging from 20 sec to 1 h and heated at 90 ºC for 3 min.
After all time points have been collected, an equal volume of formamide loading dye was added to each sample, and all samples were denatured at 90 ºC for 5 min and loaded onto a 15% polyacrylamide gel in 1x TBE buffer. After electrophoresis at 1500 V for approximately 1.5 h, the gel is then wrapped and exposed overnight on a storage phosphor screen for image capture. The phosphor screen was then scanned using GE Healthcare Typhoon Trio scanner and quantified *via* ImageQuaNT (v8.2). The resulting data was analyzed using GrafIt (v5.0.10) to determine the active fraction and rate of enzymatic turnover (under multiple turnover conditions), $k_3$, or rate of glycosidic bond cleavage, $k_{gly}$ (under single turnover conditions).

*Inductively Coupled Plasma-Mass Spectrometry*
Samples were prepared for metal analysis by inductively couple plasma-mass spectrometry (ICP-MS) as previously described.[4] Metal analysis was utilized to measure the amount of Fe and Zn ions in the purified protein sample to determine metal content of each mutant. Blanks were prepared with 45% Heparin B with buffers used in FPLC purification of each mutant and submitted in triplicate. Samples were submitted in triplicate to the UC Davis Interdisciplinary Center for ICP-MS by diluting 500 µl sample (either protein or blank) with 1.5 ml 3% nitric acid. Analysis was performed by Austin Cole to determine the role ratio of Fe and Zn to Mutyh.

*Fluorescence polarization*
To prepare the substrate for fluorescence polarization, the OG and 6'FAM-FA containing strands were annealed by pre-heating at 90 ºC for 5 min followed by an overnight annealing step at 4 ºC. DNA mixtures were prepared with final concentrations of 20 mM Tris-HCl pH 7.6, 10% glycerol, 100 mM NaCl, 1 mM EDTA, 0.1 mg/ml BSA, and 5 nM labeled duplex. The DNA mixture was aliquoted into tubes to be combined with an equal volume of enzyme. Enzyme dilutions were made at 4 ºC such that final active enzyme concentrations were between 0 to 1.5 µM. Upon addition of enzyme to DNA, tubes were incubated at 25 ºC for 30 min before loading in triplicate into a black 384-well plate (Greiner Bio-One). Fluorescence polarization was then measured on the BMG Labtech Clariostar multimode plate reader. The fluorescence polarization data was then analyzed and graphed on GrafIt(v5.0.10) to determine the $K_{1/2}$. To determine the bound fraction, the fluorescence polarization values were normalized to the negative control and plotted on GrafIt (v5.0.10) to determine the $K_{1/2}$ value for each variant.

*General cell culture*
All cell culture described herein was conducted using HEK293 cells. The wild-type HEK293FT Flp-In competent cells were provided by Dr. Wolf-Dietrich Heyer's laboratory. The matched MUTYH knockout cells were generated by CRISPR-Cas9 by laboratory member Savannah Conlon. Cells were grown at 37 ºC with 5% $CO_2$ with 4.5 g/L D-glucose Dulbecco's modified Eagle's medium (Gibco) supplemented with 10% fetal bovine serum (Gibco), 1% non-essential amino acids (Gibco), and 1% GlutMAX (Gibco). Cells were harvested for flow cytometry using 0.05% trypsin-EDTA (Gibco) and resuspended in 5% FBS in 1x PBS.

*Stable cell line generation and validation*
To generate stable cell lines, the Flp-In$^{TM}$ System from ThermoFisher was used in the MUTYH$^{-/-}$ cells. Codon optimized wild-type and variant *MUTYH* genes were delivered to these cells using the *pcDNA5$^{TM}$/FRT/TO* vector. The wild-type 521 isoform was cloned into the *pcDNA5$^{TM}$/FRT/TO* vector using Gibson assembly. Plasmids carrying each of the mutants were subsequently purchased from Twist Biosciences. The *MUTYH*-containing plasmid was co-transfected with the pOG44 recombinase containing plasmid using the Attractene transfection agent according to manufacturer's protocol. Selection was performed using a final concentration of 50 µg/ml hygromycin for two weeks.
Single colonies were identified after selection and cultured for validation using zeocin sensitivity, reverse transcriptase polymerase chain reaction (RT-PCR), and western blotting, where applicable. Zeocin sensitivity was conducted by plating cells with 0 to 600 µg/ml zeocin for two weeks. Proper recombination into the FlpIn site will disrupt the zeocin resistance gene and result in cell death even at the lowest concentration of 50 µg/ml zeocin. RNA isolation was conducted using the RNAqueous-4PCR Kit and manufacturer's protocol and subsequently quantified *via* Nanodrop. RT-PCR was conducted with the Qiagen OneStep RT-PCR kit according to the manufacturer's protocol with the corresponding primers (Table S3).

Validation *via* western blotting was conducted on all cell lines except the Gln338X truncated cell lines, as the commercially available MUTYH antibody recognizes the C-terminal domain of the protein, which is truncated in this mutant. Cells between 50-70% confluency were rinsed with 1x PBS before being harvested using RIPA buffer (150 mM NaCl, 1% Nonidet P-40, 0.5% sodium deoxycholate, 0.1% SDS, 25 mM Tris-HCl pH 7.6 and a crushed protease inhibitor tablet) and a cell scraper. Cell lysates were incubated on ice for 15 min, vortexed twice with 1 min in between, and incubated for another 15 min. After incubation, the lysates were pelleted by centrifugation at 15,000 rpm for 30 min at 4 ºC. The supernatant was removed for subsequent quantification and analysis. Supernatants were quantified using the ThermoScientific Pierce™ BCA Protein Assay Kit according to the manufacturer's protocol with the microplate procedure after diluting the supernatant 1:5 in sterile 1x PBS. Each sample was run in triplicate; 5 µl of each sample was loaded into a 96-well plate (ThermoScientific Nunclon Delta Surface) with 200 µl of the working reagent. The plate was incubated at 37 ºC for 30 min in the dark before cooling to room temperature to be measured at 562 nm on a BMG Labtech Clariostar multimode plate reader. The protein concentration of each sample was calculated using the standard curve generated alongside the samples.

Samples were loaded onto a 10% SDS-PAGE gel such that each sample contained 15 µg of total protein. After electrophoresis at 100 mA for 1-2 h, samples were transferred onto a nitrocellulose membrane using the wet transfer method. Transfer was confirmed using the reversible Ponceau S stain prior to blocking with 5% dry milk (w/v) in 1x PBS-T (1x PBS supplemented with 0.05% Tween-20) at room temperature for 1 h. After blocking, blots were incubated with the primary antibody overnight at 4 ºC. For MUTYH (Abnova 4D10), the antibody was diluted 1:240 in 1x PBS-T; for the housekeeping gene vinculin (Sigma V9131), the antibody was diluted 1:2000. After overnight incubation, the blots were washed and probed with the corresponding secondary antibody for 1 h at room temperature. Both antibodies were blotted with the anti-mouse secondary antibody conjugated to HRP at a concentration of 1:10,000 in 1x PBS-T. The blots were washed after incubation and briefly incubated with freshly mixed West Femto Max Sensitivity HRP substrate for image capture on a BioRad ChemiDoc and quantified on ImageJ.

*Plasmid purification for mammalian cellular repair assays*

A total of four plasmids are required for flow cytometry experiments. To prepare the negative and positive control plasmids, the *pUC19*, GFP OFF, and GFP ON plasmids were transformed into chemically competent DH5a and purified using a Qiagen Miniprep kit. However, the GFP OFF plasmid used to generate the OG:A mispair for use in the mammalian cell assay must be first purified using a cesium chloride gradient to produce the highest quality plasmid for nicking. Commercial miniprep and midiprep kits contain an alkyl lysis step that may damage the plasmid or hinder downstream reactions, thus the CsCl purification was shown to produce the optimal starting material for OG:A plasmid generation. This GFP OFF plasmid contained a T:A base pair in the location where the OG:A mispair would be later installed. The positive control GFP ON plasmid contained a G:C base pair in the same position.

The GFP OFF plasmid to be used for generating the OG:A plasmid required purification by using the CsCl gradient. The plasmid was first transformed into a chemically competent cell line for plasmid purification, such as DH5a or Top10. A single colony was used to inoculate a 1 L of LB supplemented with the appropriate antibiotics and grown overnight at 37 ºC and 220 rpm. Cells were then harvested by centrifugation and resuspended in 200 ml of autoclaved ice-cold STE (10 mM Tris-HCl, pH 8.0, 0.1 M NaCl, 1 mM EDTA, pH 8.0). The cells were then centrifuged at 4,100 rpm for 15 min at 4 ºC, and pellets were subsequently stored at -80 ºC before purification. To begin purification, pellets were thawed on ice before resuspension in 100 ml of ice-cold Tris-Sucrose solution (50 mM Tris-HCl pH 8.0, 10% sucrose). Next, 20 ml 10 mg/ml lysozyme and 40 ml 0.25 M EDTA were added to the suspension and incubated for 10 min on ice. Then, 40 ml 10%

SDS was added to the solution and gently stirred with a glass stir rod before addition of NaCl to a final concentration of 1 M. The slurry was incubated on ice for at least 1 h before removal of proteins by centrifugation at 12,000 rpm for 30 min at 4 ºC. The supernatant was further filtered through cheese cloth before addition of 2 volumes of 100% ethanol and incubation for 2 h. The solution was centrifuged again at 12,000 rpm for 30 min at 4 ºC, and the plasmid-containing pellet was rinsed with 1 volume of cold 70% ethanol and subsequently resuspended in 10-15 ml 1x TE buffer. Then, 1.28 volumes of 7.6 M NaI was added to the resuspended pellet and incubated for 5 min at room temperature. One volume of 100% isopropanol was added to the solution and incubated for 15 min at room temperature. The plasmid was then pelleted at 12,000 rpm for 30 min at 4 ºC and rinsed with cold 70% ethanol.

The pellet is resuspended in 3 ml of 1x TE buffer, to which 15 µl 10 mg/ml RNase A was added to incubate for 15 min at room temperature. Next, 0.1 volume 3 M NaOAc pH 5.2 and 1.0 volume isopropanol were added to the solution to incubate for 30 min at room temperature. The solution is pelleted by centrifugation at 12,000 rpm for 30 min at 4 ºC and rinsed with 1 volume of cold 70% ethanol. The pellet was air dried before resuspending in 1 ml 1x TE buffer. The plasmid was then run on a 1% agarose gel to confirm plasmid molecular weight before proceeding.

The resuspended plasmid was brought up to a final volume of 19 ml with 1x TE buffer with a final concentration of 6.2 M CsCl. The solution was brought up to 20 ml by addition of 1 ml of 10 mg/ml ethidium bromide. The solution was mixed thoroughly before being transferred into 2 self-sealing tubes for ultracentrifugation at 50,000 rpm at room temperature for a minimum of 20 h. The lower band containing unnicked plasmid was extracted under long UV illumination and purified away from the ethidium bromide using *n*-butanol saturated with 1x TE buffer until the organic phase was transparent. Six volumes of ice-cold 100% ethanol was added to the plasmid containing solution and incubated at 4 ºC overnight to precipitate the plasmid DNA. The solution was spun at 19,000 rpm for 30 min at 4 ºC, and the pellet was rinsed with 70% ethanol and spun again for 10 min. The final pellet was resuspended in 1x TE buffer and quantified by Nanodrop for subsequent experiments.

*Generation of the OG:A containing plasmid*

We used an optimized lesion-reporter plasmid reported recently to evaluate the repair of modified "OG" substrates.[5]The overall strategy was similar to that we had also reported previously, where the plasmid contains an OG:A lesion within the GFP gene and the *dsRed* transfection control gene.[6] The OG:A mispair was created within the *GFP* gene such that proper repair initiated by MUTYH would result in transcription read through and a glycine residue incorporated into the full-length protein, whereas no repair or improper repair would result in a premature stop codon and no green fluorescence.

To prepare the OG:A containing plasmid, 4 µg of CsCl purified plasmid (see Plasmid purification for mammalian cellular repair assays above) was first digested by the Nb.Bpu10I nicking enzyme for 4 h at 37 ºC. Complete nicking was confirmed *via* agarose gel prior to setting up ligation mixtures with 10 pmol phosphorylated GFP OG oligo (see Preparation of oligonucleotide substrates). Prior to adding the ligase, the mixture was heated at 90 ºC for 5 min and allowed to cool slowly to room temperature over the course of approximately 3 h. To initiate ligation, 1000 U ligase was added to the ligation mixture and incubated at room temperature for 2 h. Digestion of plasmids that did not incorporate the OG containing oligo was conducted using the AfeI restriction digest enzyme at 37 ºC overnight. The reactions were then treated with T5 exonuclease to remove all digested plasmid, leaving only the OG containing plasmid to be transfected. After installation of the OG:A mispair, the plasmid was purified and concentrated using a MACHEREY-NAGEL NucleoSpin® Gel and PCR Clean-up kit according to the manufacturer's protocol.

*Transfection and flow cytometry*
Cells were plated in a 24-well plate such that they were at 40-50% confluency at the time of transfection. Each well was transfected with a total of 400 µg total plasmid: non-fluorescent *pUC19* plasmid (negative control), GFP OFF plasmid, OG:A containing plasmid with *pUC19* carrier, or the GFP ON plasmid (positive control) using the Attractene transfection agent. The *pUC19* carrier aids in facilitating cellular uptake of the OG:A plasmid and reduces the amount of lesion containing plasmid necessary for transfection. Cells were harvested by trypsinization approximately 48 h after transfection and resuspended in 1x PBS supplemented with 5% FBS. Cells were then filtered through a 12x75mm flow cytometry tube and stored on ice prior to analysis on a Beckman Coulter CytoFLEX. The reported percent OG:A repair was determined as the fraction of transfected cells that also display green fluorescence. Additionally, repair in each cell line was normalized to the GFP ON control.

### Supporting Figures

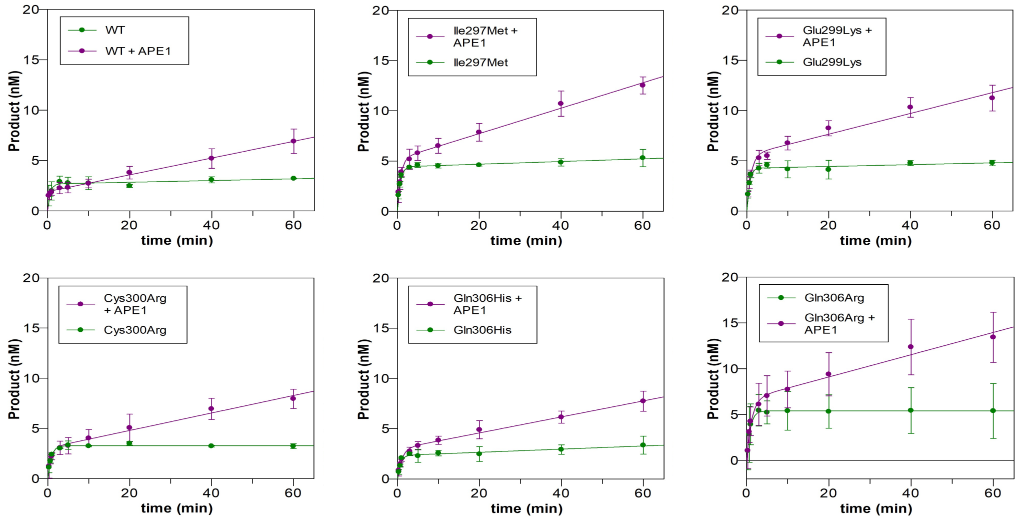


Figure S1: Stimulation of Mutyh variants by APE1

Turnover of Mutyh was assessed in the absence (green) and presence (purple) of wild type Mutyh and each purified MAP variant. In all cases, enzymatic turnover was increased in the presence of APE1.

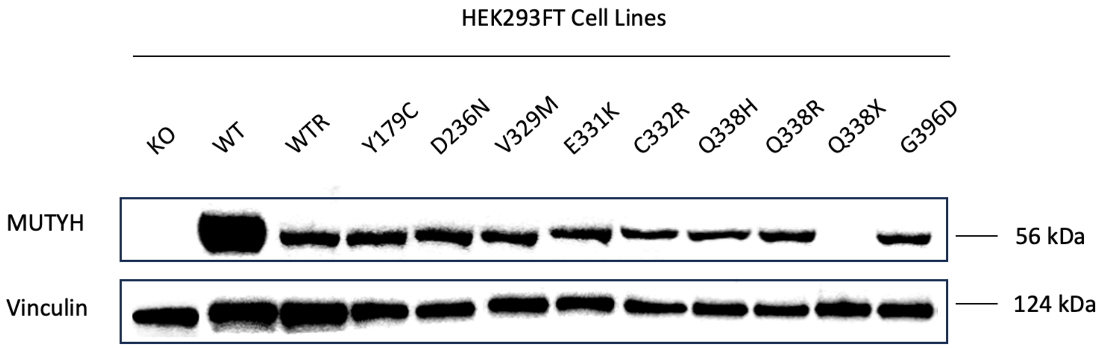


Figure S2. Western Blot Confirmation of Generated Cell Lines. See Stable cell line generation and validation in Methods.

A

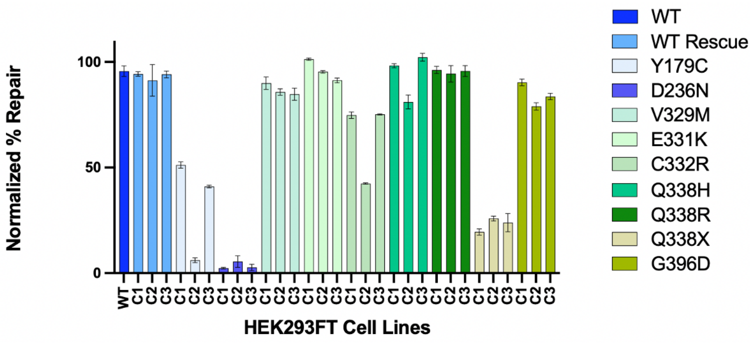


B

| | Normalized OG:A Repair (%) | | |
|---|---|---|---|
| Cell Line | Colony 1 | Colony 2 | Colony 3 |
| WT Rescue | 94 ± 1 | 91 ± 9 | 94 ± 1 |
| Y179C | 51 ± 1 | 6 ± 2 | 41± 1 |
| D236N | 5 ± 2 | 3 ± 1 | 2 ± 1 |
| V329M | 90 ± 3 | 86 ± 2 | 85 ± 3 |
| E331K | 101 ± 1 | 95 ± 1 | 91 ± 1 |
| C332R | 72 ± 8 | 42 ± 1 | 75 ± 1 |
| Q338H | 98 ± 1 | 81 ± 1 | 102 ± 1 |
| Q338R | 96 ± 2 | 94 ± 4 | 96 ± 4 |
| Q338X | 20 ± 1 | 26 ± 1 | 24 ± 2 |
| G396D | 91 ± 2 | 79 ± 1 | 84 ± 2 |

Figure S3. Normalized OG:A % Repair per Colony

A) Repair from each mutant is plotted as an average within each clone or colony. Each bar represents triplicate analysis across a single clone in any given cell line.

B) B) Data is tabulated as mean + the standard deviation after triplicate analysis per colony.

A

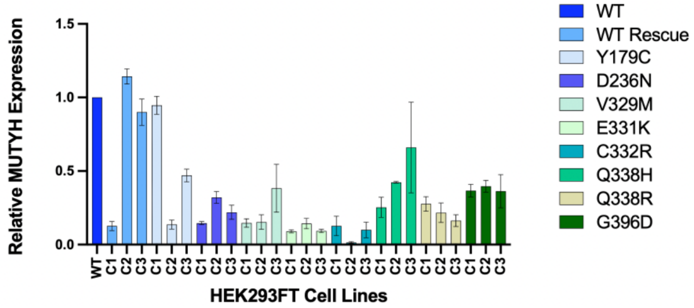


B

| | Relative MUTYH Expression | | |
|---|---|---|---|
| Cell Line | Colony 1 | Colony 2 | Colony 3 |
| WT Rescue | 0.1 ± 0.03 | 1.2 ± 0.1 | 0.9 ± 0.1 |
| Y179C | 0.9 ± 0.06 | 0.1 ± 0.03 | 0.5 ± 0.04 |
| D236N | 0.2 ± 0.05 | 0.3 ± 0.04 | 0.2 ± 0.04 |
| V329M | 0.2 ± 0.03 | 0.2 ± 0.03 | 0.4 ± 0.2 |
| E331K | 0.1 ± 0.01 | 0.1 ± 0.04 | 0.1 ± 0.01 |
| C332R | 0.1 ± 0.07 | 0.01 ± 0.006 | 0.1 ± 0.05 |
| Q338H | 0.3 ± 0.05 | 0.2 ± 0.07 | 0.2 ± 0.04 |
| Q338R | 0.3 ± 0.07 | 0.4 ± 0.006 | 0.7± 0.3 |
| G396D | 0.4 ± 0.04 | 0.4 ± 0.04 | 0.4 ± 0.1 |

Figure S4. Relative MUTYH expression in selected clones of generated cell lines

A) Average expression per colony (triplicate loading) relative to WT endogenous MUTYH expression. For each loading (triplicate), the MUTYH band was quantified as a ratio to the vinculin loading control band, and then assessed relative to WT endogenous MUTYH expression.

B) Data is tabulated as mean + the standard deviation after triplicate analysis per colony.

Table S1: Primers used to create mutants used in this study

| Mutation | Forward Primer | Reverse Primer |
|---|---|---|
| I297M | 5'-CCAGGCCGTCCTGACATGGAGGAGTGTGCTCTCAAC-3' | 5'-GTTGAGAGCACACTCCTCCATGTCAGGACGGCCTGG-3' |
| E299K | 5'-GGCCGTCCTGACATAGAGAAGTGTGCTCTCAACACTAGACAGTGC-3' | 5'-GCACTGTCTAGTGTTGAGAGCACACTTCTCTATGTCAGGACGGCC-3' |
| Q306R | 5'-GCTCTCAACACTAGACGGTGCCAGCTTTGCCTCCCTCCC-3' | 5'-GGGAGGGAGGCAAAGCTGGCACCGTCTAGTGTTGAGAGC-3' |
| Q306X | 5'-GCTCTCAACACTAGATAGTGCCAGCTTTGCCTCCCTCCC-3' | 5'-GGGAGGGAGGCAAAGCTGGCACTATCTAGTGTTGAGAGC-3' |
| Flanking | 5'-AGCTCCGTCGACTCAGCCAGGCCAAGCCTTCC-3' | 5'-TCGAGTGCGGCCGCTCACTGGGTAGTACTGTTGGGT-3' |

Table S2: Oligonucleotide sequences

| Name of oligonucleotide | Sequence |
|---|---|
| Duplex I (OG strand) | 5’- CGATCATGGAGCCAC(OG)AGCTCCCGTTACAG - 3’ |
| Duplex I (A/fA strand) | 5’ - CTGTAACGGGAGCTXGTGGCTCCATGATCG - 3’ where x = A or fA |
| GFPOG | 5’ – TGAGGCATGGAAGCGCTGACTCCCGTTGC- 3’ |

Table S3: Primers for stable cell line validation.[a]

| Primer | Sequence |
|---|---|
| Twist-MUTYHgene_Fwd | 5’-GGCCATGGAGCTGGGTGCTACAG-3’ |
| Twist-MUTYHgene_rev | 5’-CCAGCAAGATTTGAGCGCCCAGGG-3’ |
| primer 243_fwd | 5’-GGACCTACCATGGAGAAGACG-3’ |
| primer 1090_rev | 5’-CTGCACCAGCAGAATTTGG-3’ |

[a]Twist-MUTYHgene primers were used for cell lines generated using the codon optimized MUTYH variant genes ordered from Twist Biosciences. The wild-type non-codon optimized cell line was confirmed using primer 243_fwd and primer 1090_rev.
